# Toll-like receptor 5 as a novel receptor for fungal zymosan

**DOI:** 10.1101/2021.12.23.473960

**Authors:** Stefanie Reuter, Kristina Herold, Jana Domrös, Ralf Mrowka

## Abstract

Microbial pathogens carry specific structural patterns which were termed pathogen-associated molecular patterns (PAMPs). Toll-like receptors (TLRs) as key elements for the recognition of microbial pathogens are necessary for the activation of innate immune pathways. TLRs are activated by binding PAMPs of bacteria, viruses and fungi and initiate a signaling pathway resulting in the activation of transcription factors which modulate the production of various proinflammatory cytokines. It is not fully clear in detail which microbial pattern is recognized by which TLR. Here we show for the first time that TLR5 is a strong receptor for the yeast particle zymosan. We have generated stable human cell lines with combinations of TLR2 and TLR5 knock in/knock out together with stable nuclear factor kappaB (NF-κB) luciferase reporters. We found that both receptors TLR5 and TLR2 lead to an independent activation of the NF-κB pathway when simulated with zymosan. Our results demonstrate that TLR5 is a receptor for the fungal particle zymosan in addition to bacterial fragments like flagellin. Distinct cytokine patterns might suggest that TLR5 is potentially important for the differentiation in the recognition of the specific type of the foreign microorganisms and in the specific host defense response.

## Introduction

Microbial pathogens like bacteria, viruses and fungi carry a huge number of microbial structures named pathogen-associated molecular patterns (PAMPs) which initiate the activation of numerous innate immune pathways by binding to specific pattern recognition receptors (PRRs). The family of Toll-like receptors (TLRs) belongs to PRRs and consists of type I transmembrane proteins. The structure of all known human TLRs is characterized by a conserved cytoplasmic region that is termed Toll/IL-1R (TIR) domain, an extracellular domain and a transmembrane domain. The TIR domain contains three highly homologues regions which are necessary for downstream signaling whereas the extracellular domain of TLRs comprises leucin-rich repeat (LRR) motifs and is responsible for recognizing molecular patterns from microbial components^1,2^. TLRs are expressed on macrophages, dendritic cells, B cells and neutrophils. Moreover, non-immune cells like fibroblast cells, epithelial cells and keratinocytes comprise TLRs^2,3^.

Currently 10 members of the TLR family are known in humans. TLR1, TLR2, TLR4, TLR5 and TLR6 are anchored on the cell surface and are involved in the recognition of external microbial components such as lipopeptides, lipopolysaccharide and flagellin^4,5^. TLR2 recognizes predominantly structures which are associated with the cell wall of Gram-positive bacteria, bacterial lipoproteins and zymosan from fungi and mycobacterial components^6,7^. TLR2 is known as receptor to discriminate yeast and Gram-positive bacteria from Gram-negative bacteria^8^. TLR5 is characterized as receptor for flagellated bacteria by recognizing flagellin^9,10^. A successful ligand binding and activation of the receptor requires a homo or hetero dimerization of single TLRs^11^. Activated receptor complexes initiate a downstream signaling cascade which triggers the activation and translocation of the transcription factors nuclear factor-κB (NF-κB), interferon regulatory factor 3/7 (IRF3/7) and activator protein-1 (AP-1) into the nucleus. This initiates the production of inflammatory cytokines, type I interferon and chemokines^12,13^. After stimulation of TLRs with their specific ligand, each TLR shows a different cytokine expression profile. Especially macrophages are involved in cytokine production after the activation of TLRs^4^.

In this work we investigated the role of TLR5 in fungal ligand recognition by the usage of a luciferase reporter gene assay for TLR-induced signaling via NF-κB. Therefor we generated firefly luciferase HEK293 reporter cells stably expressing single TLRs or combinations of TLRs and cofactors. TLR activation and signaling induces the activation and translocation of the transcription factor NF-κB. NF-κB binds to a designed sensitive promoter element consisting of three different consensus sequences for the NF-κB subunits p65 and p50 and activates the firefly luciferase generation.

## Results

### Activation of TLR2 and TLR5 through the fungal cell wall component zymosan

TLRs are pattern recognition receptors that recognize a wide range of molecular patterns. To study TLR activation by the fungal cell wall component zymosan and other ligands we performed cellular reporter assays, based on firefly luciferase HEK293 reporter cells. The firefly luciferase reporter gene is controlled by an artificial promoter containing three different consensus sequences for the NF-kB subunits p65 and p50. Each consensus sequenz is repeated six to seven times (supplementary figure S1). Binding of NF-κB to the promoter element induces bioluminescence that is measured in living cells. HEK293 reporter cells endogenously not express TLR2, but TLR5. Endogenously expressed TLR5 was knocked out by the use of Cas9 targeted to exon 6 of the TLR5 encoding gene. To perform cellular reporter assays we used HEK293 reporter cells stably and endogenously expressing TLR5 referred to as HEK-TLR5 reporter cells and HEK293 reporter cells stably expressing TLR2 and its cofactor CD14 referred to as HEK-TLR2-CD14 reporter cells. Furthermore, in HEK-TLR2-CD14 reporter cells endogenously expressed TLR5 was knocked out by the use of Cas9 targeted to exon 6 of the TLR5 encoding gene.

To characterize the cell lines HEK293 reporter cells, HEK-TLR5 reporter cells and HEK-TLR2-CD14 reporter cells were treated with the TLR2 ligand FSL-1, the TLR5 ligand flagellin and the heat inactivated Gram-positive bacteria *Staphylococcus aureus* (*S. aureus*) and the Gram-negative bacteria *Salmonella typhimurium* (*S. typhimurium*). HEK-TLR5 reporter cells were activated by the specific ligand flagellin (**Figure 1a**) and *S. typhimurium* (**Figure 1b**). HEK-TLR2-CD14 reporter cells were activated by the specific ligand FSL-1 (**Figure 1a**) and *S. typhimurium* as well as *S. aureus* (**Figure 1b**). In HEK293 reporter cells, which were stimulated as control cell line, the luciferase activity did not increase after the treatment with any of the added stimulants (**Figure 1a and 1b**). Flagellin and *S. typhimurium* are able to activate TLR5. FSL-1, *S. typhimurium* and *S. aureus* are able to activate TLR2.

**Figure 1.**
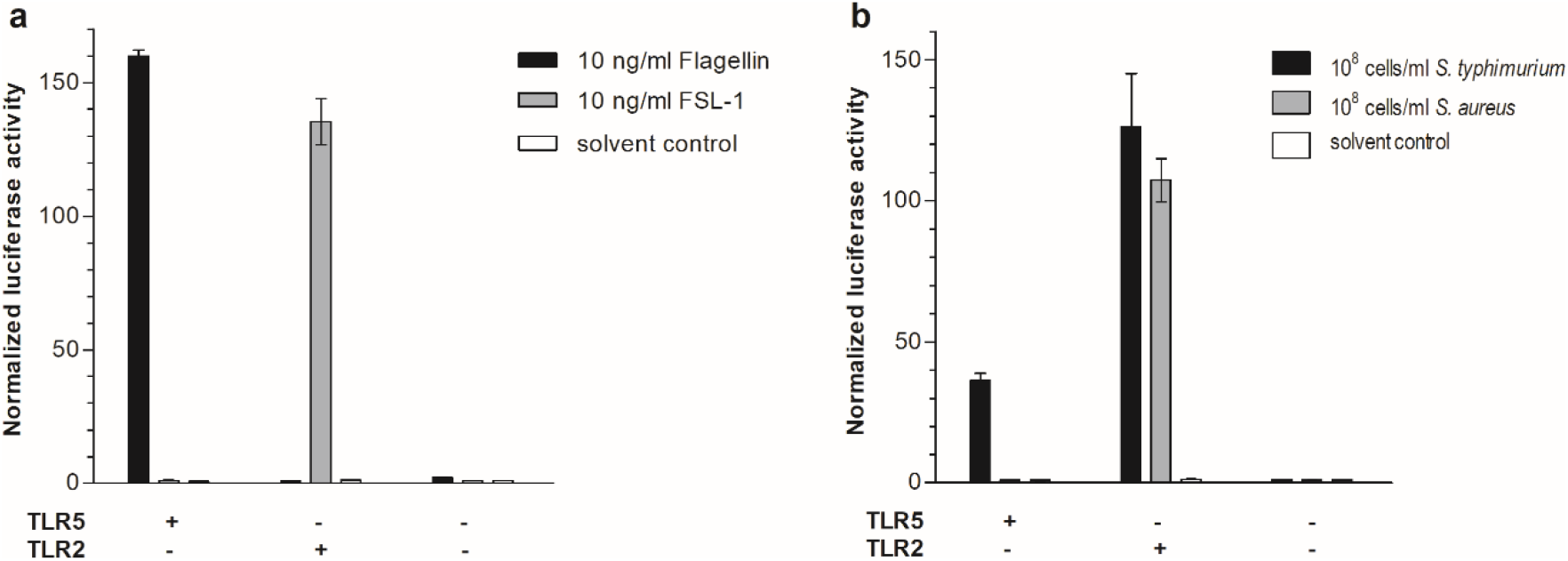
Activation of HEK293 reporter cells after treatment with TLR2 and TLR5 ligands. HEK-TLR5 reporter cells stably and endogenously expressing TLR5, HEK-TLR2-CD14 reporter cells stably expressing TLR2 and with TLR5 knock out and HEK293 reporter cells without endogenously TLR2 expression and with TLR5 knock out were treated with the TLR2 ligand FSL-1, the TLR5 ligand flagellin and the heat inactivated Gram-positive bacteria *S. aureus* and the Gram-negative bacteria *S. typhimurium* (n=5). Data represents activation 10h after stimulation. (a) In HEK-TLR5 reporter cells the luciferase activity increased after the treatment with 10 ng/ml Flagellin, whereas the treatment with 10 ng/ml FSL-1 did not increase the luciferase activity. In HEK-TLR2-CD14 reporter cells treatment with 10 ng/ml FSL-1 leads to an increased luciferase activity, whereas treatment with 10 ng/ml Flagellin did not. In HEK293 reporter cells luciferase activity did not increase after the treatment with neither Flagellin nor FSL-1. (b) In HEK-TLR5 reporter cells the luciferase activity increased after the treatment with 10^8^ cells/ml *S. typhimurium,* whereas the treatment with 10^8^ cells/ml *S. aureus* did not increase the luciferase activity. In HEK-TLR2-CD14 reporter cells treatment with 10^8^ cells/ml *S. typhimurium* and 10^8^ cells/ml *S. aureus* leads to an increased luciferase activity. In HEK293 reporter cells luciferase activity did not increase after the treatment with neither *S. typhimurium* nor *S. aureus*. Error bars typify standard deviation (two-way ANOVA: p<0,0001, receptor: p<0,0001, treatment p<0,0001).

Zymosan is a complex molecule consisting of glucan, mannan, protein and lipoprotein components and is a cell wall preparation of the yeast *Saccharomyces cerevisiae* (*S. cerevisiae*)^14,15^. Zymosan is recognized by TLR2 in cooperation with CD14^16,17^. Furthermore, the C-type lectin receptor Dectin-1 collaborates with TLR2 during the recognition of zymosan^18^. HEK293 reporter cells, HEK-TLR5 reporter cells and HEK-TLR2-CD14 reporter cells were treated with zymosan in four different concentrations (10 μg/ml, 5 μg/ml, 2,5 μg/ml and 1 μg/ml). The treatment with zymosan leads to a concentration dependent increase of the luciferase activity in HEK-TLR5 reporter cells (**Figure 2a**) and in HEK-TLR2-CD14 reporter cells (**Figure 2b**). HEK293 reporter cells were stimulated as a control cell line and did not respond to the treatment with zymosan (**Figure 2c**). Zymosan activates its specific receptor TLR2. In our cellular reporter assay we found, that zymosan can activate TLR5, too. Hot alkali-treated zymosan, referred to as depleted zymosan lost the ability to activate TLR2 in former studies^19^. We used this depleted zymosan in stimulation experiments, but did not observe any activation in HEK-TLR5 or HEK-TLR2-CD14 expressing cells (**Figure 2d, 2e, 2f**).

**Figure 2.**
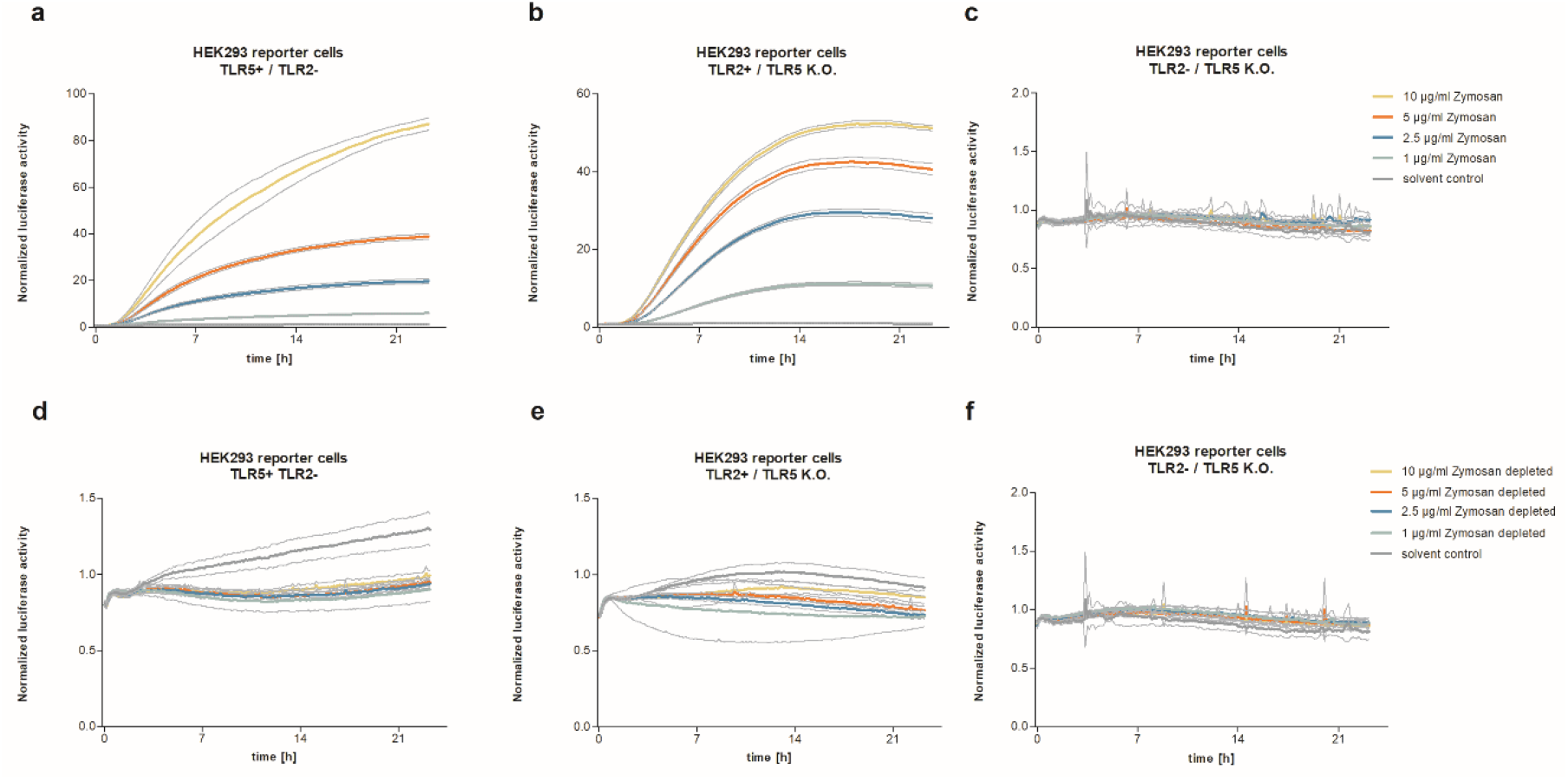
Activation patterns of HEK293 reporter cells after treatment with zymosan and “zymosan depleted” substances depending on the TLR status. HEK293 reporter cells were treated in the dose range of 0-10 μg/ml zymosan and “zymosan depleted” (n=5). (a) In HEK-TLR5 reporter cells stably and endogenously expressing TLR5 luciferase activity increased in a dose and time dependent manner after treatment with zymosan. (b) In HEK-TLR2-CD14 reporter cells stably expressing TLR2 and with TLR5 knock out luciferase activity increased in a dose and time dependent manner after treatment with zymosan, too. (c) In HEK293 reporter cells with TLR5 knock out and without endogenous TLR2 expression luciferase activity did not increase after the treatment with zymosan. The substance “zymosan depleted” has no TLR activation properties. After treatment of HEK-TLR5 reporter cells stably and endogenously expressing TLR5 (d) and HEK-TLR2-CD14 reporter cells stably expressing TLR2 and with TLR5 knock out (e) luciferase activity did not increase. (f) HEK293 reporter cells without endogenously TLR2 expression and with TLR5 knock out expression did not respond to the added zymosan depleted, too. Error lines show the standard deviation.

For a deeper understanding of the potential binding site of TLR5 on zymosan, TLR5 and TLR2 antagonists were added to stimulation experiments (**Figure 3**). A soluble TLR5 receptor (hTLR5-Fc), a recombinant mouse TLR5 Fc chimera and a chimeric monoclonal antibody specific for human TLR5 (anti-hTLR5-IgA) function as competitive inhibitors to block TLR5 activation. A chimeric monoclonal antibody specific for human TLR2 (anti-hTLR2-IgA) was used to neutralize the biological activity of TLR2.

**Figure 3.**
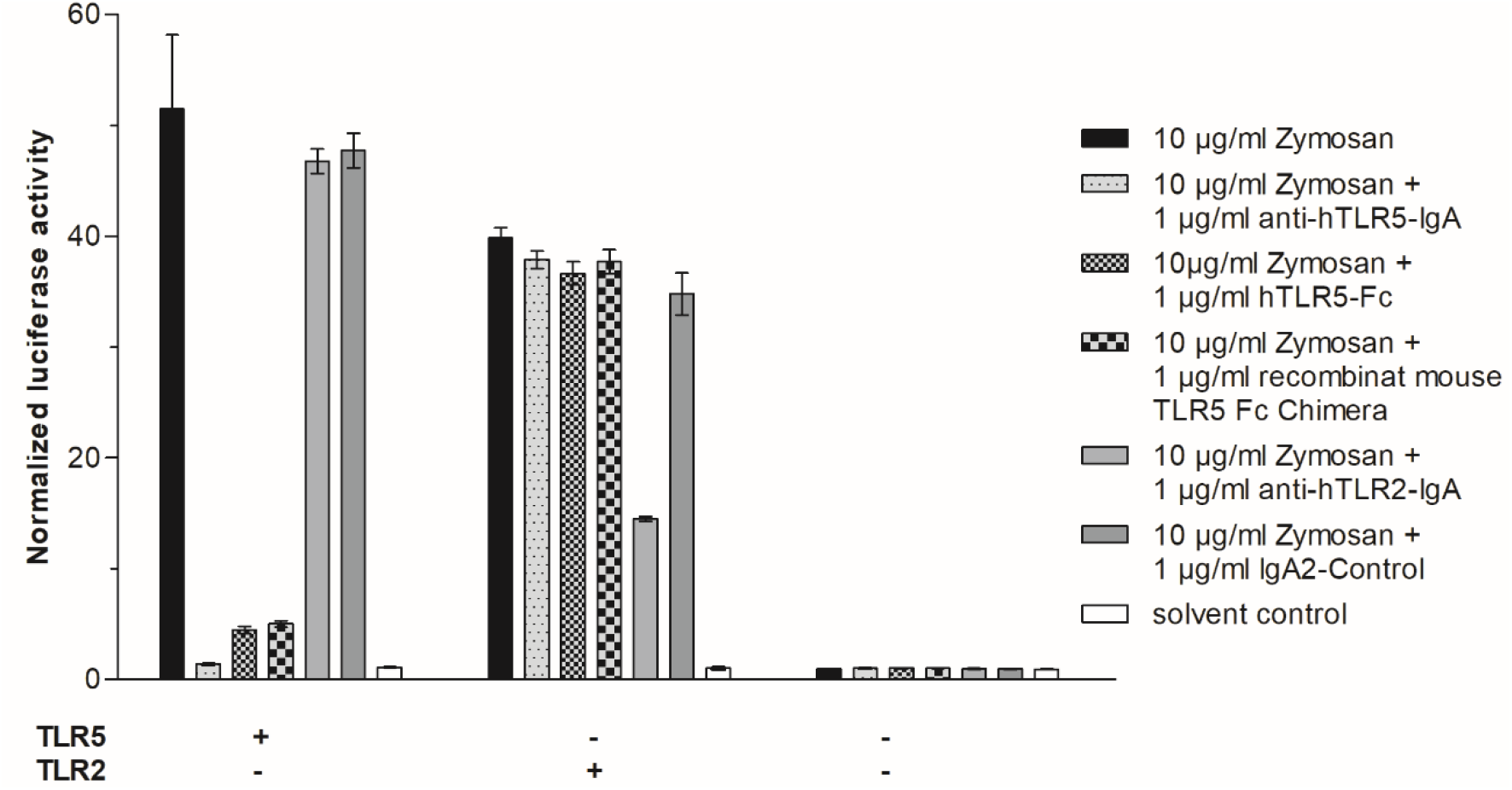
Inhibition of TLR activation by TLR antagonists. HEK-TLR5 reporter cells stably and endogenously expressing TLR5, HEK-TLR2-CD14 reporter cells stably expressing TLR2 and with TLR5 knock out and HEK293 reporter cells without endogenously TLR2 expression and with TLR5 knock out were treated with 10 μg/ml zymosan in combination with 1 μg/ml of different competitive TLR5 antagonists, a TLR2 antagonist and a IgA2-control. Data represents reaction of the cells 10h after stimulation (n=5). Zymosan induced increase of the luciferase activity is inhibited by TLR5 antagonists in HEK-TLR5 reporter cells whereas the TLR2 antagonist anti-hTLR2-IgA did not decrease zymosan induced luciferase activation in HEK-TLR5 reporter cells. Zymosan induced increase of the luciferase activation in HEK-TLR2-CD14 cells is decreased by anti-hTLR2-IgA but not by the different TLR5 antagonists. In HEK293 reporter cells luciferase activity did not increase after treatment with zymosan or zymosan in combination with TLR antagonists. Error bars typify standard deviation (two-way ANOVA: p<0,0001, receptor: p<0,0001, treatment p<0,0001).

Zymosan induced increase of luciferase activity in HEK-TLR5 reporter cells was inhibited by an anti-TLR5-IgA antibody and by human and mouse soluble TLR5 ectodomains but not by an anti-TLR2 IgA antibody. In contrast zymosan induced activation of the luciferase was inhibited by the anti-TLR2 IgA antibody but not by antagonists targeted to TLR5.

Since the soluble ectodomain of mouse TLR5 was able to inhibit zymosan-induced TLR5 activation, HEK293 reporter cells with TLR5 knock out were transiently transfected with human and mouse TLR5-encoding plasmids and stimulated by addition of zymosan, flagellin and TLR5 and TLR2 antagonists (**Figure 4**). As seen in human TLR5 stimulation, zymosan induced the increase of luciferase activity in HEK293 reporter cells which were transiently transfected with mouse TLR5 (**Figure 4c**). Mouse TLR5 activation was inhibit by human TLR5 antagonists and the soluble ectodomain of mouse TLR5 (**Figure 4d**).

**Figure 4.**
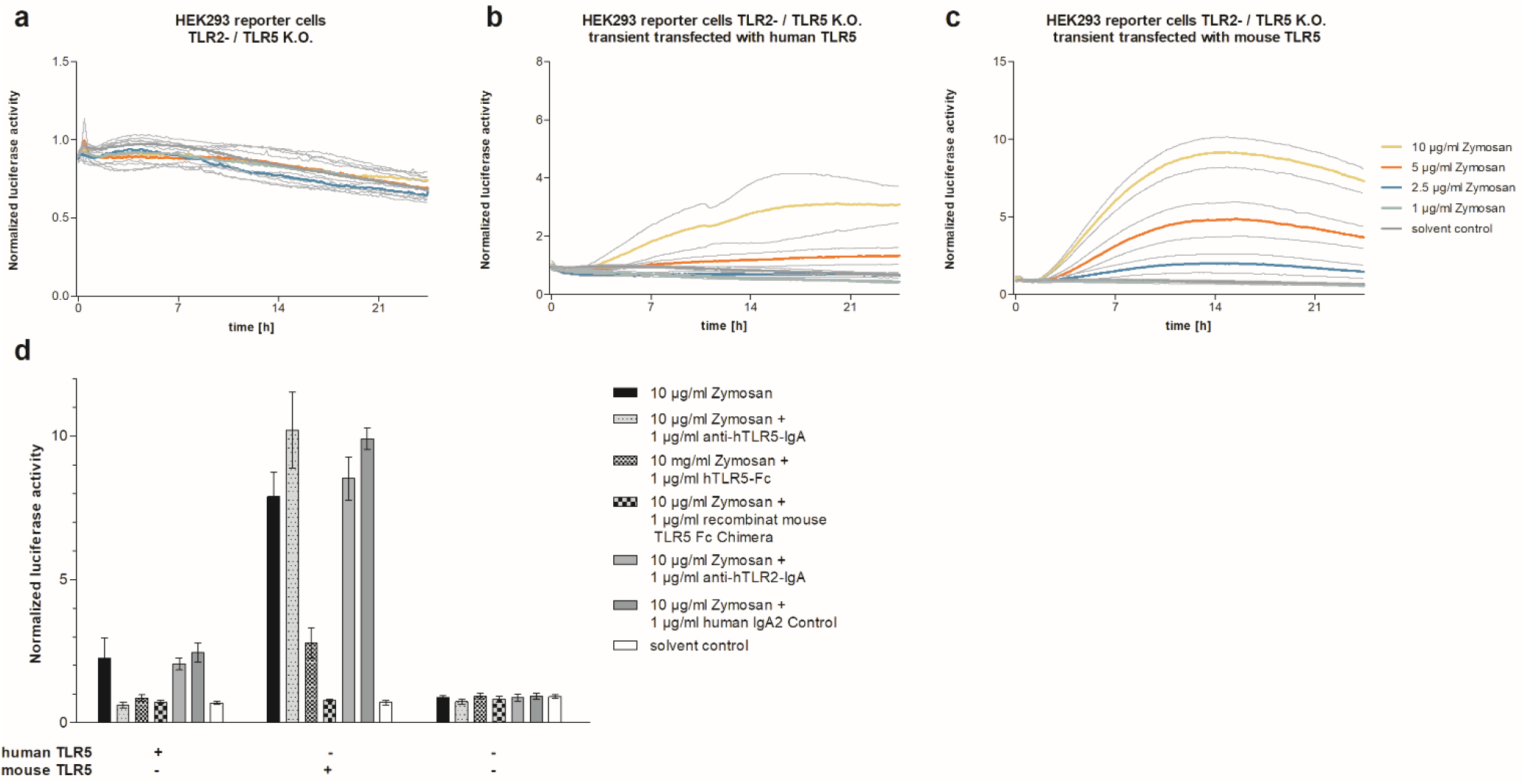
Zymosan shows also activation of mouse TLR5. HEK293 reporter cells without endogenous TLR2 expression and TLR5 knock out were transient transfected with human TLR5 or mouse TLR5. Transient transfected cells and HEK293 reporter cells were treated with the indicated doses of zymosan (n=5). (a) In HEK293 reporter cells without endogenous TLR2 expression and TLR5 knock out luciferase activity did not increase. (b) In HEK293 reporter cells which are transient transfected with human TLR5 luciferase activity increases according to zymosan concentration. (c) In HEK293 reporter cells which are transient transfected with mouse TLR5 luciferase activity increases according to zymosan concentration on a higher level compared to human TLR5. (d) Zymosan induced increase of luciferase activity in HEK293 reporter cells with human TLR5 is inhibited by TLR5 antagonists anti-hTLR5-IgA, hTLR5-Fc and recombinant mouse TLR5 Fc Chimera. Zymosan induced increase of luciferase activity in HEK293 reporter cells with mouse TLR5 is only inhibited by recombinant mouse TLR5 Fc Chimera. Error bars typify standard deviation (two-way ANOVA: p<0,0001, receptor: p<0,0001, treatment p<0,0001).

### Zymosan induced the production of proinflammatory cytokines TNFα, Interleukin 8 (IL-8) and CXC chemokine-ligand 1 (CXCL1)

Flagellin induced TLR5 activation triggers the production and release of proinflammatory cytokines like CC chemokines CXC chemokines and tumor necrosis factors via the NF-κB - dependent signaling pathway^4^. To investigate cytokine production by zymosan stimulation, HEK-TLR5 reporter cells were stimulated with zymosan and the supernatant was tested for the presence of cytokines and chemokines via R&D Proteome Profiler Cytokine Array. We detected IL-8 and CXCL1 in the supernatant of HEK-TLR5 reporter cells. In contrast flagellin-induced TLR5 activation leads to the production of IL-8, CXCL1, CCL2 and GM-CSF (**Figure 5**). HEK-TLR2-CD14 reporter cells were incubated with zymosan and FSL-1 respectively. IL-8 and CXCL1 were found in the according supernatant, whereas FSL-1 induced the production of IL-8, CXCL1 and CCL2.

**Figure 5.**
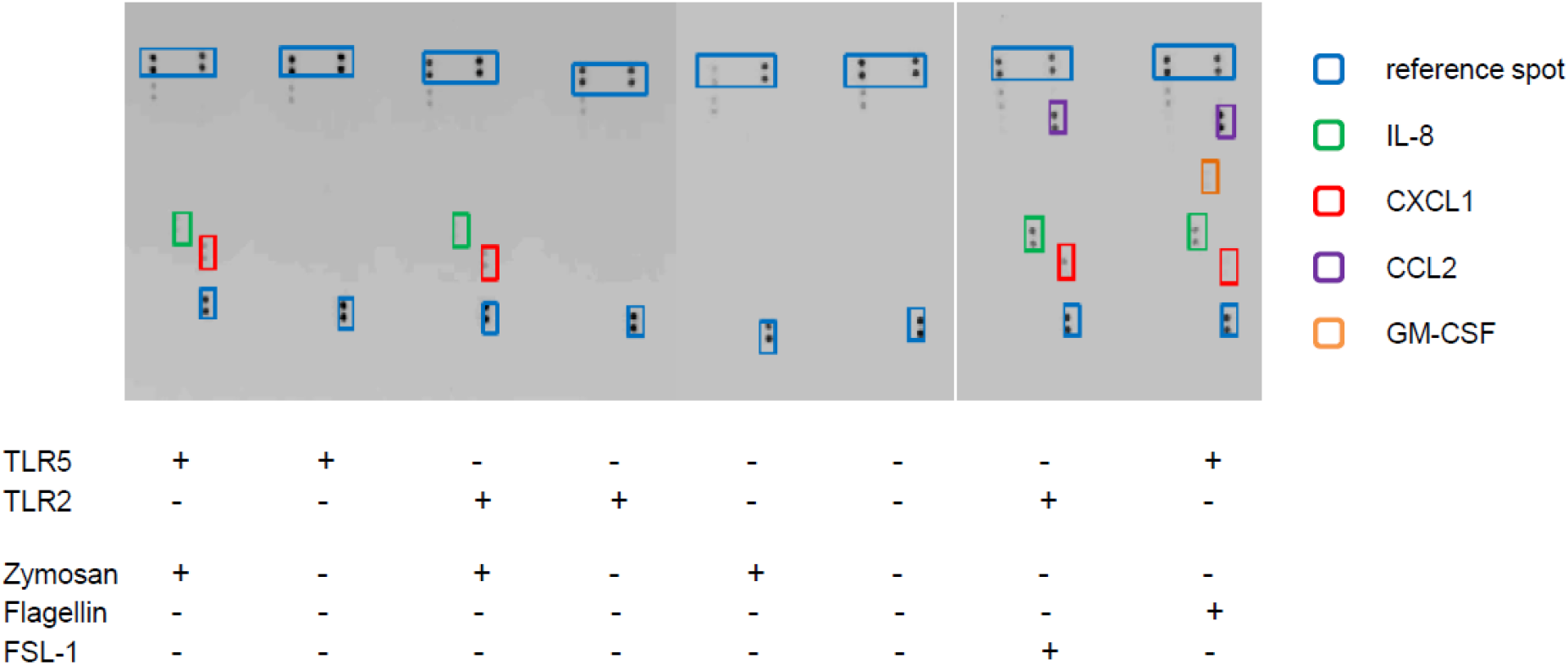
Cytokine patterns upon stimulation with TLR agonists. HEK-TLR5 reporter cells and HEK-TLR2-CD14 reporter cells were stimulated with zymosan and cytokine production was measured. After treatment with zymosan the cytokines IL-8 (green box) and CXCL1 (red box) was detected. Unstimulated HEK-TLR5 reporter cells and HEK-TLR2-CD14 reporter cells and reporter cells without TLR5 and TLR2 expression did not show the production of cytokines. After treatment of HEK-TLR2-CD14 reporter cells the cytokines IL-8 (green box), CXCL1 (red box) and CCL2 (violet box) were detected. After treatment of HEK-TLR5 reporter cells the cytokines IL-8 (green box), CXCL1 (red box), CCL2 (violet box) and GM-CSF (orange box) were detected. Blue boxes represent reference spots.

## Discussion

TLRs are evolutionary highly conserved pattern recognition receptors (PRRs) that recognize a broad range of molecular patterns associated with pathogens (PAMPs) but also endogenous molecular patterns that are released under certain circumstances like apoptosis or tissue necrosis^5^. The recognition of fungal PAMPS involves lots of PRRs, among them are TLR2 and TLR6. Underhill et al. found first that TLR2 is involved in phagocytosis of zymosan leading to the production of TNFα^8^. Sato et al. show that TLR2 directly binds to zymosan^16^. Finberg et al. initially described zymosan as TLR2 ligand through transfection experiments in HEK293 cells. The induction of TLR2 signaling by zymosan requires the cofactor CD14^17^. Heterodimers of TLR2 and TLR6 induce cytokine production upon recognition of different fungal pathogens including zymosan^20^.

Our study revealed TLR5 as a novel receptor for the fungal cell wall component zymosan. So far TLR5 is known to be the exclusive receptor for bacterial flagellin. It is located on the outer cell membrane and highly expressed on dendritic cells of the lamina propria of the small intestine where it is involved in B cell differentiation to IgA producing plasma cells^21^.

We performed cellular reporter assays, based on firefly luciferase HEK293 reporter cells stably expressing single TLRs or combinations of TLRs and cofactors. TLR activation and signaling induces the enzymatic activity of the firefly luciferase through its NF-κB sensitive promoter element consisting of three different repetitive consensus sequences for NF-κB. TLR5 expressing cells and cells expressing the combination of TLR2 and CD14 were generated by stable transfection. Furthermore, endogenously expressed TLR5 was knocked out by Cas9 targeted to exon 6 of the TLR5 encoding gene. The luciferase activity was measured before and after adding stimulants to the culture medium.

The fungal cell wall component zymosan is an activator of TLR5 signaling in a dose-dependent manner (**Figure 2a**) compared to flagellin and flagellin-bearing Gram-negative *Salmonella typhimurium* (*S. typhimurium*) bacterial cells. HEK reporter cells without TLR5 expression did not respond to zymosan (**Figure 2c**) whereas TLR2-CD14 reporter cells were activated by zymosan as well (**Figure 2b**). Ikeda et al. demonstrated that chemical treatment of zymosan with organic solvents like chloroform-methanol removed the ability of zymosan to activate TLR2 but not Dectin-1^19^, another important PRR recognizing fungal zymosan^22,23^. Such chemical treatment removes lipid-like structures that probably build the molecular structure that is essential to activate TLR2. We observed a similar effect in the cellular stimulation assays where TLR5 activation was abolished by depleted zymosan (**Figure 2d, 2e, 2f**). Probably, TLR5 recognizes lipid-like structures, too. Inhibition assays with TLR5 and TLR2 antagonists showed that a soluble human and mouse TLR5 ectodomain and an antibody against human TLR5 were able to decrease the zymosan-induced TLR5 activation but not zymosan-induced activation of TLR2-CD14 (**Figure 3**). An antibody against TLR2 inhibited zymosan-induced TLR2 activation but not TLR5 activation. TLR5 and TLR2 recognize zymosan by binding to different molecular structures of zymosan. Since the soluble ectodomain of mouse TLR5 is also able to inhibit zymosan-induced TLR5 signaling we transiently transfected the whole mouse TLR5 receptor into HEK luciferase reporter cells without endogenous TLR5 and TLR2 expression. Zymosan also activated mouse TLR5 signaling in our assay system (**Figure 4c**).

Signaling of TLRs leads to the translocation of the transcription factor NF-κB from the cytoplasma into the nucleus where it binds to promotors of cytokine encoding genes for instance. We analyzed the supernatant of zymosan-stimulated HEK-TLR5 reporter cells and found IL-8 and CXCL1 in contrast to the supernatant of flagellin-induced TLR5 activation, where IL-8, CXCL1, CCL2 and GM-CSF were found (**Figure 5**). Zymosan induced the release of IL-8 and CXCL1 in HEK-TLR2-CD14 luciferase reporter cells, too. Whereas the TLR2 specific ligand FSL-1 induced the release of IL-8, CXCL1 and CCL2 in HEK-TLR2-CD14 luciferase reporter cells. Treatment of both TLR2 and TLR5 with zymosan leads to the release of the cytokins IL-8 and CXCL1 while treatment with the specific TLR2 and TLR5 ligands shows a different cytokine pattern. TLR5 seems to be another receptor supporting antifungal innate immune reactions.

## Methods

### Plasmids for cell line generation

Lentiviral packaging plasmids pMDLg/pRRE (Addgene plasmid #12251; http://n2t.net/addgene:12251; RRID:Addgene_12251)^24^, pRSV-Rev (Addgene plasmid # 12253; http://n2t.net/addgene:12253; RRID:Addgene_12253)^24^ and pMD2.G (Addgene plasmid # 12259; http://n2t.net/addgene:122539; RRID:Addgene_12259) were a gift from Didier Trono and were obtained via Addgene.

### Cloning of the NF-κB responsive promoter and luciferase reporter gene

NF-κB responsive promoter contains three different binding motifs for the p65 and p50 subunit of NF-κB (5’-GGGGACTTTCC-3’, 5’-GGGGATTCCC-3’, 5’-GGGGATTTCC-3’)^25^,^26^. The 220 bp promoter construct was flanked by restriction enzyme sites for EcoNI (5’) and HindIII (3’) and synthesized by ATG:biosynthetics. Additional to the NF-κB binding motifs NF-κB response element and a TATA-like sequence from pNFκB-Luc vector (Clontech Cat. No. 631904) were added to the Firefly luciferase gene and introduced into pCDH-CMV-EF1-Puro.

### Luciferase reporter cell line generation by lentiviral transduction

The generation of HEK luciferase reporter cells has already been described in detail elsewhere^27^. In brief, stable integration of the luciferase reporter gene and its NF-κB responsive promoter was achieved by lentiviral transduction of Flp-In^™^-293 cells.

Lentivirus-containing supernatant was produced by calcium phosphate transfection of pCDH-CMV-EF1-Puro, containing the NF-κB responsive promoter and the firefly luciferase, and the packaging plasmids pMDLg/pRRE, pRSV-Rev and pMD2.G into HEK293T cells. Supernatant was collected 48, 72 and 96 hours post transfection, sterile filtered and immediately frozen at −80°C. Flp-In^™^-293 cells (Thermo Fisher Scientific) were transduced 3 times with lentivirus containing supernatant and selected with 1 μg/ml Puromycin (Invivogen, San Diego, USA). Surviving cells were expanded and referred to as HEK293 luciferase reporter cells.

### Cloning of TLR and cofactor combinations

Combinations of TLRs were cloned into pcDNA5/FRT (Invitrogen, Thermo Fisher Scientific, Waltham, USA). TLR2 was cut from pcDNA3-TLR2-YFP (pcDNA3-TLR2-YFP was a gift from Doug Golenbock (Addgene plasmid # 13016; http://n2t.net/addgene:13016; RRID:Addgene_13016)) by digestion with KpnI and XhoI and integrated into pcDNA5/FRT (Invitrogen), cut with KpnI and PspOMI, together with CD14, digested with HindIII and NotI from pcDNA3-CD14 (pcDNA3-CD14 was a gift from Doug Golenbock (Addgene plasmid # 13645; http://n2t.net/addgene:13645; RRID:Addgene_13645). Both open reading frames were connected by the TLR2-CD14 adapter oligonucleotide containing a T2A sequence. The complete open reading frame consists of TLR2-T2A-CD14 under control of the CMV promoter in pcDNA5/FRT. TLR5-CFP was digested from pcDNA3-TLR5-CFP (pcDNA3-TLR5-CFP was a gift from Doug Golenbock (Addgene plasmid # 13019; http://n2t.net/addgene:13019; RRID:Addgene_13019)) with XhoI and BamHI and ligated into pcDNA5/FRT, digested with XhoI and BamHI.

### Stable integration of TLRs by Flp-In^™^-System

5*10^6^ HEK293 luciferase reporter cells were seeded into T75 cell culture flasks in 10 ml DMEM with 10% FCS and transfected with 2 μg of pcDNA/FRT containing cloned TLR combinations and 18 μg of pOG44 and 40 μl Roti ^®^ -Fect (Carl Roth, Karlsruhe, Germany) according to the manufacturer’s protocol. 24 hours post transfection cells were washed with 10 ml DPBS (Thermo Fisher Scientific) and 10 ml DMEM containing 10% FCS, 1% Pen/Strep (10,000 U/ml) was added. 48 hours after transfection cells were splitted in a ratio of 1:4 and 300 μg/ml Hygromycin B (Thermo Fisher Scientific) was added to the cell culture medium for antibiotic selection of stably transfected cells. After 24 hours medium was changed to DMEM with 100 μg/ml Hygromycin B and selection was continued until cell foci appeared. Cell lines were referred to as HEK293-TLR2-CD14 luciferase reporter cells and HEK293-TLR5 luciferase reporter cells.

### TLR5 Knock out with CRISPR/Cas9

For Cas9-induced TLR5 knock-out, the online guide design tool **Fehler! Linkreferenz ungültig.** was used to design the following oligos: TLR5-Guide-forward CCGGGACCACCTGGACCTTCTCCT and TLR5-Guide-reverse AAACAGGAGAAGGTCCAGGTGGTC. Both guides were cloned into pGuide-it-tdTomato (TaKaRa Bio Europe) according to the manufacturer’s protocol.

1.5 million HEK293-TLR2-CD14 luciferase reporter cells and HEK293 luciferase reporter cells were seeded per well of a six well plate (Greiner Bio-One) in 2 ml DMEM supplemented with 10% FBS and 1% Penicillin/ Streptomycin (Pen/Strep 10,000 U/ml) and incubated for 12-16 hours in a humidified incubator at 37°C and 5% CO2. Cells were transfected with 3 μg pGuide-it-tdTomato with inserted TLR5-specific guides and 12 μl Lipofectamine 2000 transfection reagent (Invitrogen) per well. 24 hours post transfection cells were washed with DPBS, trypsinized and collected in 1ml DPBS. Cells positive for red fluorescence were sorted with a BD FACSAria^™^ Fusion and single cell deposition into 96-well cell culture plates was performed. Single cells were grown in 300 μl conditionized DMEM supplemented with 10% FBS, 1% Pen/Strep and 10 μM StemMACS Y27632 (Miltenyi). 24 hours later medium was changed to DMEM supplemented with 10% FBS and 1% Pen/ Strep. Cells were expanded and analyzed for stimulation by Flagellin as described above. Non-responding cells were characterized by amplifiying a product of TLR5 with the following primers: TLR5_forward-2 TGTCTGTATCTGCCAACAGC and TLR5_reverse-2 AAGGAGAGCGTTTTCCCTTGT. The product of 800 bp was sent for sequencing to Eurofins Genomics, Germany.

### Luciferase reporter assay

50000 HEK293 luciferase reporter cells were seeded per well of a 96 well plate (Greiner Bio-one, Frickenhausen, Germany) in 145 μl DMEM containing 4.5 g/l glucose (Thermo Fisher Scientific), 10% FCS, 1% Pen/Strep, 1 mM L-Glutamin and 250 μM Luziferin D (Promega, Madison, USA). Cells were incubated 12-16 hours in a humidified incubator at 37°C and 5% CO_2_ to adhere to the bottom of the plate. The plate was sealed prior to the measurement in a TopCount^®^NXT (PerkinElmer, Waltham, USA) microplate luminescence counter at 37°C. The cells were run without stimulation for 2 hours (pre-run) and stimulated afterwards by adding 10 μl of stimulant, diluted in cell culture medium. Luminescence counts were normalized to the mean luminescence count of the pre-run period. TLR agonists FSL-I, Flagellin, Zymosan, heat-killed *Staphylococcus aureus* (*S. aureus*) and *Salmonella typhimurium* (*S. typhimurium*) as well as TLR antagonists anti-TLR5-IgA, anti-TLR5-Fc human, anti-TLR2-IgA and IgA control antibody were purchased from Invivogen. Anti-TLR5-Fc mouse was purchased from R&D Systems, Inc.

### R&D Proteome Profiler Cytokine Array

3 million TLR5-KO-HEK-NF-κB luciferase reporter cells with stable expression of TLR2-CD14 and TLR5 expressing HEK-NF-κB luciferase reporter cells and unchanged HEK-NF-κB luciferase reporter cells were seeded per well of a six well plate (Greiner Bio-One) in 2 ml DMEM supplemented with 10% FBS and 1% Penicillin/ Streptomycin (Pen/Strep 10,000 U/ml) and incubated for 24 hours in a humidified incubator at 37°C and 5% CO_2_. The medium was changed to cell culture medium containing 1 μg/ml zymosan (InvivoGen Europe, France), 10 ng/ml FSL1 (InvivoGen), and 50 ng/ml flagellin (InvivoGen), respectively and plates were incubated for 24 hours. Medium without stimulant was used as unstimulated control. Cell supernatant was collected and 1 ml was used for Proteome Profiler^™^ Human Cytokine Array (ARY005B, R&D Systems, Inc., USA) following the manufacturer’s protocol. Chemiluminescence detection was performed at a G:BOX iChemiXL (Synoptics, UK) with exposure times from 1 to 30 minutes.

## Acknowledgements

Funding: This work was funded by BMBF FKZ 13GW0245B FungoSens to RM. We would like to thank Silke Nossmann and Gesine Hauschild for the technical assistance.

## Author contribution

SR, KH and RM wrote and edited the manuscript. JD performed TLR5 ligand assays. SR and KH generated the cell lines, performed the luciferase assays and cytokine arrays. Genomic editing was performed by SR. Data analysis and interpretations were performed by all authors. The study was designed by RM and SR.

## Additional Information

All authors declare not conflict of interest.

---

**Supplementary Figure S1.**
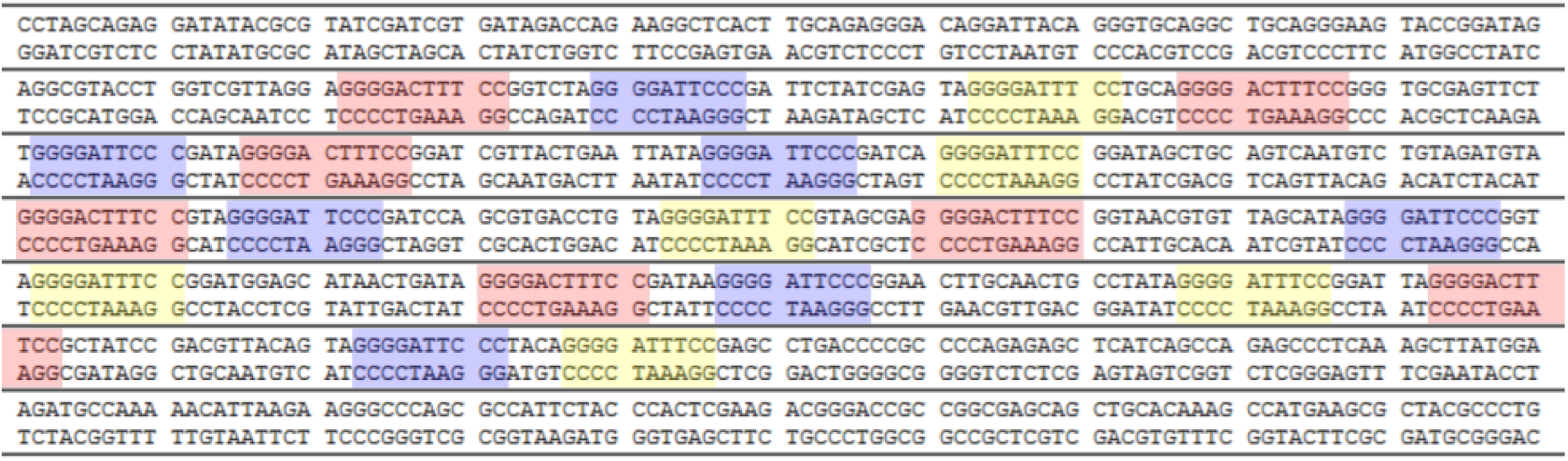
The NF-κB sensitive promotor region consists of the three different NF-κB consensus sequences 5’- GGGGACTTTCC- 3’, 5’- GGGGATTCCC- 3’ and 5’- GGGGATTTCC- 3’. These NF-κB consensus sequences are specific binding motives for the NF-κB subunits p65 and p50.

